# DFAST_QC: Quality Assessment and Taxonomic Identification Tool for Prokaryotic Genomes

**DOI:** 10.1101/2024.07.22.604526

**Authors:** Mohamed Elmanzalawi, Takatomo Fujisawa, Hiroshi Mori, Yasukazu Nakamura, Yasuhiro Tanizawa

## Abstract

**Motivation:** Accurate taxonomic assignments of genomic data are crucial across various biological databases. With a rapid increase in submitted genomes in recent years, ensuring precise classification is important to maintain database integrity. Mislabeled genomes can confuse researchers, hinder analyses, and produce false results. Therefore, there is a critical need for computationally efficient tools that ensure accurate taxonomic classification for data to be deposited into genomic databases.

**Results:** Here we introduce DFAST_QC, a quality control and taxonomic classification tool of prokaryotic genomes based on NCBI and GTDB taxonomies. We benchmarked DFAST_QC’s performance against NCBI taxonomy assignments, showing high consistency with them. Our results demonstrate that DFAST_QC achieves high consistency to NCBI taxonomy classification.

**Availability and implementation:** DFAST_QC is implemented in Python and is available both as a web service (https://dfast.ddbj.nig.ac.jp/dqc) and as a stand-alone command line tool. The source code is available under the GPLv3 license at: https://github.com/nigyta/dfast_qc, and the conda package is also available from Bioconda. The data and scripts used for the benchmarking process are publicly available on GitHub (https://github.com/Mohamed-Elmanzalawi/DFAST_QC_Benchmark).

**Supplementary information:** Supplementary data are available at Bioinformatics online.

## 1 Introduction

Public genome databases are fundamental to biological research, where ensuring accurate metadata and high-quality sequences supports open data practices and facilitates collaborative research efforts. However, taxonomically mislabeled genomes within the databases can cause confusion or lead to scientifically inaccurate results when referenced or reused in other researches (Bagheri et al., 2020; Goudey et al., 2022).

To ensure accurate taxonomic labeling, the National Center for Biotechnology Information (NCBI) has used Average Nucleotide Identity (ANI) analysis since 2018 to verify prokaryotic genomes in GenBank (Ciufo *et al*. 2018). ANI is a method that compares the genetic similarity between two genomes by calculating the mean identity of the homologous regions from the pairwise alignment between two genomes, with a threshold of 95% ANI commonly accepted to distinguish species (Goris *et al*. 2007). Within the International Nucleotide Sequence Database Collaboration (INSDC), taxonomic information is organized based on NCBI Taxonomy to maintain the consistency and interoperability of the organism names (Schoch *et al*. 2020). As such, the NCBI Taxonomy serves as a standard, although not authoritative, resource for nomenclature and classification.

DFAST_QC has been developed as a validation tool for the DNA Data Bank of Japan (DDBJ), a member of INSDC, to ensure accurate taxonomic assignment and quality of prokaryotic genomes submitted to the database. It is incorporated into the DFAST web service, a genome annotation and data submission pipeline for DDBJ (Tanizawa, Fujisawa and Nakamura 2018), but also functions as a standalone tool that can run on a local machine. DFAST_QC performs quick taxonomic identification based on NCBI Taxonomy using Mash (Ondov *et al*. 2016) similarity estimation and Skani (Shaw and Yu 2023) for accurate ANI calculation. It also assesses genome completeness and contamination using CheckM (Parks *et al*. 2015). It can optionally query against representative genomes in the GTDB Taxonomy (Parks *et al*. 2022). In this paper, we will present the features of DFAST_QC, focusing on taxonomic identification.

## 2 Materials and methods

### 2.1 Workflow of DFAST_QC

DFAST_QC performs taxonomy checks using a two-step approach to reduce running time while maintaining accuracy. The required input is a simple FASTA file. Initially, the FASTA file undergoes genomic distance calculation using the MASH sketch file generated from reference genome sequences. In the second step, Skani is used to create a sketch file for these genomes, resulting in a manageable sketch file size and increased process speed. Then, ANI is calculated between the query genome and the selected reference genomes to determine taxonomic assignment, applying species-specific ANI thresholds when available, or using a default threshold of 95%. For the quality assessment, CheckM is employed to assess the completeness and contamination percentage of the query genome. The marker set for CheckM is automatically determined based on the result of the taxonomy check or can be specified manually. Finally, the genome size is checked to ensure it falls within the expected range. When specified as an option, species identification is performed based on GTDB Taxonomy by searching against its representative genomes. A detailed workflow figure can be found in Supplementary Fig. S2 and an example use case of DFAST_QC can be found in Supplementary Text S1.

### 2.2 Preparation of the reference data

DFAST_QC utilizes two primary sources of reference data: NCBI Datasets and GTDB. They can be accessed and managed using Python scripts included in the software package. A detailed figure for data preparation can be found in Supplementary Fig. S2.

#### Reference Data for NCBI Taxonomy

DFAST_QC first retrieves metadata on genomic assemblies from GenBank (assembly_summary_genbank.txt) and identifies type strains (type genomes) from this dataset. Subsequently, it filters out genomes excluded from RefSeq or identified as misidentified type genomes, using criteria defined in the ‘assembly type category’, ‘excluded from RefSeq’, and ‘taxonomy check status’ columns within the NCBI-provided file (ANI_report_prokaryotes.txt). Following this, DFAST_QC proceeds to download the filtered genomes. Afterward, it creates an SQL database that integrates information from both ”ANI_report_prokaryotes.txt” and ”assembly_summary_genbank.txt”. Finally, DFAST_QC utilizes MASH to sketch the entire genomes and generate a consolidated sketch file. To identify the ANI threshold for each species and indistinguishable groups, DFAST_QC retrieves the “prokaryote ANI indistinguishable groups.txt” and “prokaryote ANI species specific_threshold.txt” files from NCBI.

#### Reference data for GTDB Taxonomy

DFAST_QC downloads representative genomes and their metadata file from GTDB, and then it establishes a dedicated SQL database optimized for searches within GTDB. Finally, it generates another sketch file using a similar methodology as described earlier.

### 2.3 Benchmarking

To evaluate the performance of DFAST_QC, we conducted a series of comparative benchmarks. The reference data for DFAST_QC and benchmarking datasets were prepared on June 26, 2024. This reference data included 22,171 type genomes obtained from the NCBI Assembly Database and 113,104 representative genomes from the Genome Taxonomy Database (GTDB) release 220. Two benchmarking datasets, A and B, were prepared to evaluate the accuracy of species assignment based on the NCBI and GTDB taxonomies, respectively. Dataset A comprises 5,184 of the latest non-type genomes, with one genome per species randomly selected from the NCBI GenBank. Based on NCBI’s quality control, we excluded species lacking available type genomes, those with failed or inconclusive taxonomy checks, and those deemed suppressed, or contaminated. Dataset B comprises 10,000 randomly selected metagenome-assembled genomes (MAGs) from the GEMs dataset (Nayfach et al., 2021).

Genomes in both Datasets A and B were processed by DFAST_QC (ver. 1.0.0) with default settings using a single CPU for each run. Genomes in Dataset B were also processed using the classify workflow (classify_wf) of GTDB-Tk (Chaumeil *et al*. 2022) version 2.4.0 with reference data release 220.

## 3 Results and discussion

The accuracy of species assignments based on NCBI Taxonomy was assessed using Dataset A, which comprises a collection of randomly selected non-type genomes from GenBank. We compared the species names assigned by DFAST_QC using the top ANI hit to the species name as labeled for the submitted genome in GenBank (declared species). The results of this comparison are summarized in Table 1.

**Table 1.**
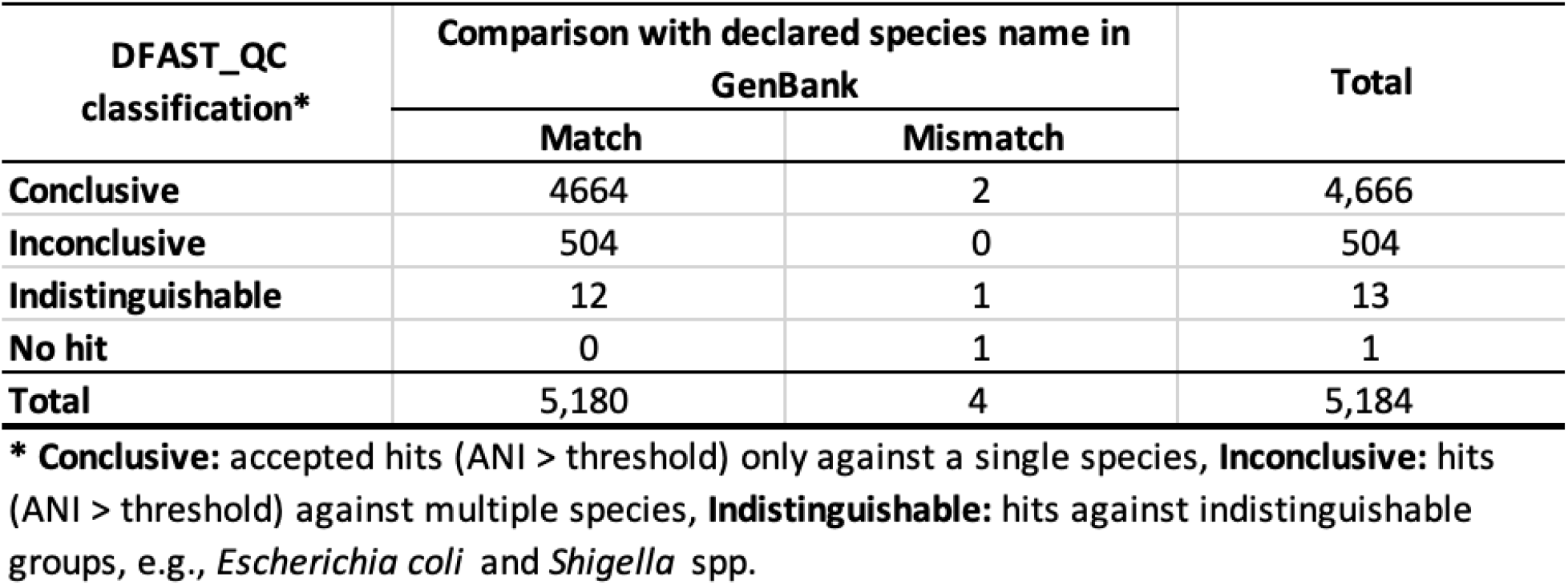
DFAST_QC results for 5184 non-type genomes from GenBank.

| DFAST_QC<br>classification* | Comparison with declared species name in<br>GenBank |  | Total |
| --- | --- | --- | --- |
|  | Match | Mismatch |  |
| Conclusive | 4664 | 2 | 4,666 |
| Inconclusive | 504 | 0 | 504 |
| Indistinguishable | 12 | 1 | 13 |
| No hit | 0 | 1 | 1 |
| Total | 5,180 | 4 | 5,184 |

\* **Conclusive:** accepted hits (ANI > threshold) only against a single species, **Inconclusive:** hits (ANI > threshold) against multiple species, **Indistinguishable:** hits against indistinguishable groups, e.g., *Escherichia coli* and *Shigella* spp.

Out of the 5,184 cases compared, the species name assigned by DFAST_QC matched the declared species in GenBank in 5,180 cases (99.9%), including 504 cases with accepted ANI hits (ANI > threshold) against multiple species and 12 cases that fell into indistinguishable species groups. The four mismatch cases were likely due to the mislabeling of the declared species or the inconsistency of the current taxonomic system (see Supplementary Text S2). For many of the Inconclusive cases, the declared species name were found in the top hit (416 cases) or in the second or lower hits with an ANI value slightly below the threshold (88 cases), indicating that they belonged to species difficult to differentiate by ANI or might be an outlier within the species.

The benchmarking using Dataset B showed high consistency with the results from GTDB-Tk at the species-level identification (Supplementary Text S3).

The benchmark demonstrates DFAST_QC’s accuracy in species identification when a reference type genome is available. However, for species lacking a sequenced type genome, DFAST_QC cannot definitively assign species. Highlighting this limitation is crucial, as NCBI’s report (prokaryote_without_type_assembly.txt) indicates that over 5,000 species currently lack sequenced type genomes. Fortunately, this situation is improving thanks to large sequencing projects like the Global Catalogue of Microorganisms (GCM) 10K type strain genome sequencing project (Wu and Ma 2019), and the growing recommendation to deposit genome sequences alongside new taxon descriptions (Riesco and Trujillo 2024). Also, as exemplified in the four mismatch cases of Dataset A (Supplementary Text S2), the functionality to search against GTDB representative genomes can serve as a complement to the results of taxonomy checks, particularly when reference genomes are not available.

Unlike other genome-based identification tools such as TYGS (Meier-Kolthoff and Göker 2019) and GTDB-Tk, DFAST_QC’s results are limited to species-level identification, with no phylogenetic inference at higher taxonomic ranks. This is because our focus is more on the correct assignment of organism names for genomes to be submitted to public sequence databases. Due to its simplicity, DFAST_QC operates on machines with limited computational resources. In fact, it requires less than 2GB of memory and can typically complete taxonomy identification within 30 seconds. Additionally, we provide a minimal set of prebuilt reference data containing only sketch files and metadata (<1.5GB in size). Although this approach results in extra execution time since reference genomes required for ANI calculation are retrieved during runtime, it eliminates the need to prepare a full set of reference data (>100GB), making installation on local machines easier.

## 4 Conclusion

DFAST_QC is a tool designed for quality and taxonomy check of prokaryotic genomes, utilizing NCBI and GTDB taxonomies for species identification. It is integrated into the web service of DDBJ’s genome annotation and submission pipeline, DFAST, featuring a user-friendly interface for researchers unfamiliar with the command-line operation. In addition, it is also available as a stand-alone software, which enables rigorous validation of genomes on a local machine before submission to public databases. It employs compact reference data and requires low computational resources. This comprehensive functionality reinforces its importance in maintaining the accuracy and reliability of genomic data across scientific research.

## Acknowledgments

Computations were partially performed on the NIG supercomputer at ROIS National Institute of Genetics. We thank Masato Suzuki and Masaki Shintani for their suggestions for identifying pathogens. We also thank Manabu Ishii for testing the software with the large-scale dataset.

