## Supplementary Document for "DFAST_QC: Quality Assessment and Taxonomic Identification Tool for Prokaryotic Genomes"

---

---

### Text S1. Example Use Case

DFAST_QC is an open-source package written in Python, available as both source code and a web version. It was designed to be user-friendly, allowing users to generate accurate and easily interpreted results with minimal effort. To demonstrate the practicality and versatility of DFAST_QC, we present a use case involving the publicly available genome of *Paucilactobacillus hokkaidonensis* (Tanizawa *et al.* 2015) on NCBI.

#### Text S1.1 Using the command line

DFAST_QC can be downloaded from our GitHub repository at https://github.com/nigyta/dfast_qc. Additionally, it is also available on Bioconda(Grüning *et al.* 2018). After downloading the reference data, users can execute the following command: "`dfast_qc -i Paucilactobacillus_hokkaidonensis.fa -o example --enable_gtdb`".

DFAST_QC must be provided with a FASTA file to run. This file path is passed to the script through the "`–i`" argument. The results are stored in a directory named after the "`–o`" argument. If omitted, the results will be saved in a directory named 'OUT'. By default, DFAST_QC performs taxonomy classification on the Genbank database. The optional parameter "`--enable_gtdb`" allows users to perform taxonomy classification using the GTDB database. To search exclusively in the GTDB database, the optional command "`—-disable_tc`" can be used to disable the GenBank database search. Another optional argument is "`–a`", which is an integer and sets the minimal ANI threshold for genomes to be included in the analysis. This filter is useful when the user aims to identify highly related genomes. The default value is set to 95%.

#### Text S1.2 Using the web version

To provide convenient and rapid access to DFAST_QC for researchers preferring a graphical user interface, we have developed a web server at https://dfast.ddbj.nig.ac.jp/dqc/submit/. Users can upload their files directly and receive results promptly. Additionally, they can specify the database for searching and set the rank and taxon parameters for CheckM to expedite results; if unspecified, these parameters are inferred automatically. Users can enter their e-mail address and a job name, to receive an e-mail after the job is completed and identify different submissions easily. The result will be deleted 30 days after the last visit.

#### Figure S1.DFAST_QC web interface:

The figure shows the main sections for file upload, parameter settings, and user input fields.

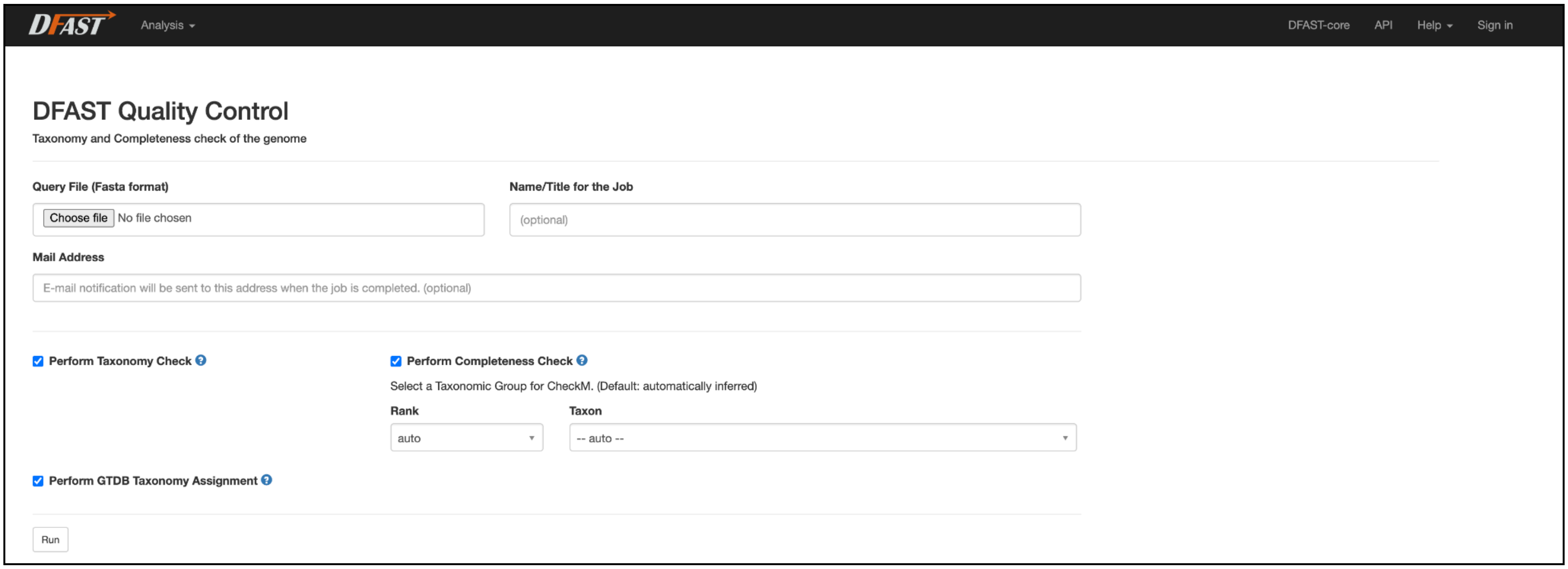

### Text S2. GenBank benchmark analysis

The 4 genomes resulting in mismatch raise concerns about potential mislabeling within the databases or issues in the current taxonomic system. Two genomes marked as conclusive in DFAST_QC, GCA_027152945.1 (*Lactobacillus gasseri*) and GCA_002354875.1 (*Actinosynnema pretiosum*), both exhibited inconsistencies.
GCA_027152945.1 is labeled as '*Lactobacillus gasseri*' in GenBank, but it was identified as '*Lactobacillus paragasseri*' with an ANI value of 98.35% by DFAST_QC. *L. paragasseri* was recently proposed as a new species, having been separated from *L. gasseri* based on ANI (Tanizawa *et al.* 2018). It was also reported that many of the genomes labeled as *L. gasseri* in the public databases belonged to *L. paragasseri* (Ene *et al.* 2022). Considering these situations, GCA_027152945.1 should correctly be labeled as '*L. paragasseri*'.

GCA_002354875.1 is identified as '*Actinosynnema pretiosum'* by the Taxonomy check in NCBI based on the identity against GCA_013387285.1 (*A. pretiosum* subsp. *auranticum* DSM $44131^{T}$). However, GCA_013387285.1 is commented as "failed to match other type_strains on its species" by NCBI's check with 93.89% ANI against GCA_024171695.1 (A. *pretiosum* subsp. *pretiosum* DSM $44132^{T}$) while showing the ANI value of 95.95% against GCA_000023245.1 (*Actinosynnema mirum*). This implies that *A. pretiosum* subsp. *auranticum* Hasegawa et al. 1983 should be treated as a later heterotypic synonym of *A. mirum* Hasegawa et al. 1978. Since the "Inconclusive" status is assigned to GCA_013387285.1 by NCBI's check, we excluded it from our reference data. This is why our result, in which GCA_002354875.1 was identified as '*A. mirum*', was incongruent with that of NCBI's check. Notably, both GCA_002354875.1 and GCA_013387285.1 are elevated into a species-level cluster named *'Actinosynnema auranticum'* in GTDB Taxonomy r220, forming a clade distinct from *A. pretiosum* and *A. mirum*. (https://www.ncbi.nlm.nih.gov/datasets/genome/GCA_002354875.1/, https://www.ncbi.nlm.nih.gov/datasets/genome/GCA_013387285.1/, https://gtdb.ecogenomic.org/species?id=Actinosynnema%20auranticum accessed on 2024/7/1).

The third case GCA_013997415.1 (*Shigella dysenteriae*) fell within an indistinguishable group according to NCBI's "prokaryote_ANI_indistinguishable_groups.txt". So it is expected to have difficult strain-level identification. However, it is worth mentioning that, since we limited our Mash top hit results to 10, it was not included in our analysis. When we increased the threshold, it appeared with ANI (97.5%), although it was lower than its ANI species-specific threshold (99.2%).

The fourth mismatch GCA_000576125.1 had no accepted hit in DFAST_QC. We found it to be a similar case as GCA_002354875.1. GCA_000576125.1 is identified as '*Zymomonas mobilis subsp. mobilis'* by NCBI's check using GCA_000175255.2 (*Zymomonas mobilis* subsp. *mobilis* ATCC 10988$^{T}$) as a reference, but GCA_000175255.2 is assigned with an "Inconclusive" status because it shows low ANI (80.41%) against the type genome of another subspecies (GCA_006539385.1, *Zymomonas mobilis* subsp. *pomaceae* NBRC 13757$^{T}$). Therefore, GCA_000175255.2 is not included in our reference data. The two subspecies should ideally be reclassified into two distinct species. In fact, in GTDB Taxonomy, they are placed in two different species-level clades. Accordingly, GCA_000576125.1 was classified as '*Zymomonas mobilis*' based on the DFAST_QC result against GTDB representative genomes. (https://www.ncbi.nlm.nih.gov/datasets/genome/GCA_000576125.1/ , https://www.ncbi.nlm.nih.gov/datasets/genome/GCA_000175255.2/, accessed on 2024/7/1).

### Text S3. GTDB-Tk benchmark

GTDBtk(Chaumeil *et al.* 2020), a standalone tool based on the Genome Taxonomy Database (GTDB), offers higher-rank classification for uncultured microbes using relative evolutionary divergence (RED) by estimating the evolutionary distance between genomes. However, its application is limited by substantial computational demand, large reference data requirements, and incompatibility with the NCBI taxonomy. This presents a limitation for researchers who need to submit data to public databases.

species-level taxonomy classification based on GTDB Taxonomy was evaluated by comparing the results from DFAST_QC and GTDB-Tk (Table S2). According to the GTDB-Tk classification, Dataset B (10,000 MAGs from GEMs) consisted of 9,390 bacterial genomes and 610 archaeal genomes.

**Table S2.GTDB-TK benchmark results:**

Comparison of Species level classification for 10000 MAGs from GEMs between DFAST_QC and GTDB-TK.

| DFAST_QC species classification | GTDB-TK species classification | | Total |
|---|---|---|---|
| | Assigned | Unassigned | |
| Assigned | Identical / 7056<br>Mismatch / 68 | 33 | 7,157 |
| Unassigned | 2 | 2,841 | 2,843 |
| Total | 7,126 | 2,874 | 10,000 |

**"Assigned":** Genomic sequences classified into specific taxonomic species with ANI > 95%.
**"Unassigned":** Genomic sequences that could not be classified.

DFAST_QC successfully assigned species names to 7,124 genomes, with 7,056 (99%) of these classifications matching those made by GTDB-Tk. The remaining 68 genomes exhibited mismatches in species classification, primarily due to multiple candidate genomes with average nucleotide identity (ANI) values around 95% or closely similar ANI values, highlighting the challenges in precise species identification.

One major difference between DFAST_QC and GTDB-Tk is a higher-rank taxonomic classification. GTDB-Tk can assign higher-rank classification based on RED even if no species-level classification was available, whereas DFAST_QC cannot give higher-rank information when no accepted ANI hits were found. However, hits against close relatives may give some indication. Also, GTDB-Tk can handle multiple genomes in a batch, while DFAST_QC can only process one genome per run. This drawback can be alleviated by invoking multiple DFAST_QC processes in parallel, as it can run with a small memory footprint (<4GB) and in a short time, typically less than 1 minute for the taxonomy check.

**Figure S2.DFAST_QC workflow figure.**

The figure outlines the DFAST_QC workflow, detailing steps for ANI calculation, sketch file creation, and quality assessment. Key processes include the use of MASH, Skani, and CheckM evaluation.

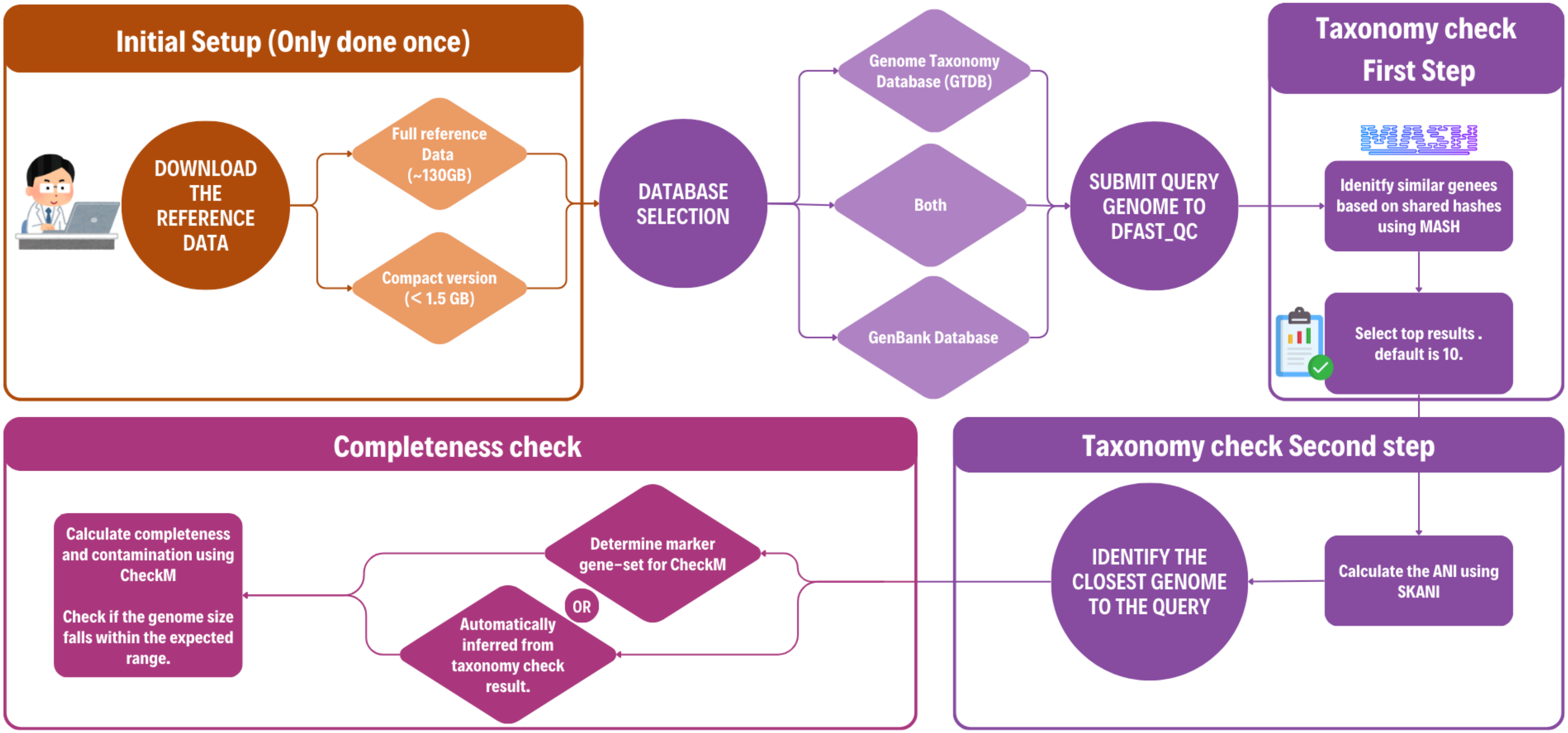

**Figure S3.Reference data preparation.**

The figure depicts the steps for preparing reference data for DFAST_QC, including metadata retrieval, genome filtering, SQL database creation, and genome sketching for both NCBI and GTDB taxonomies.

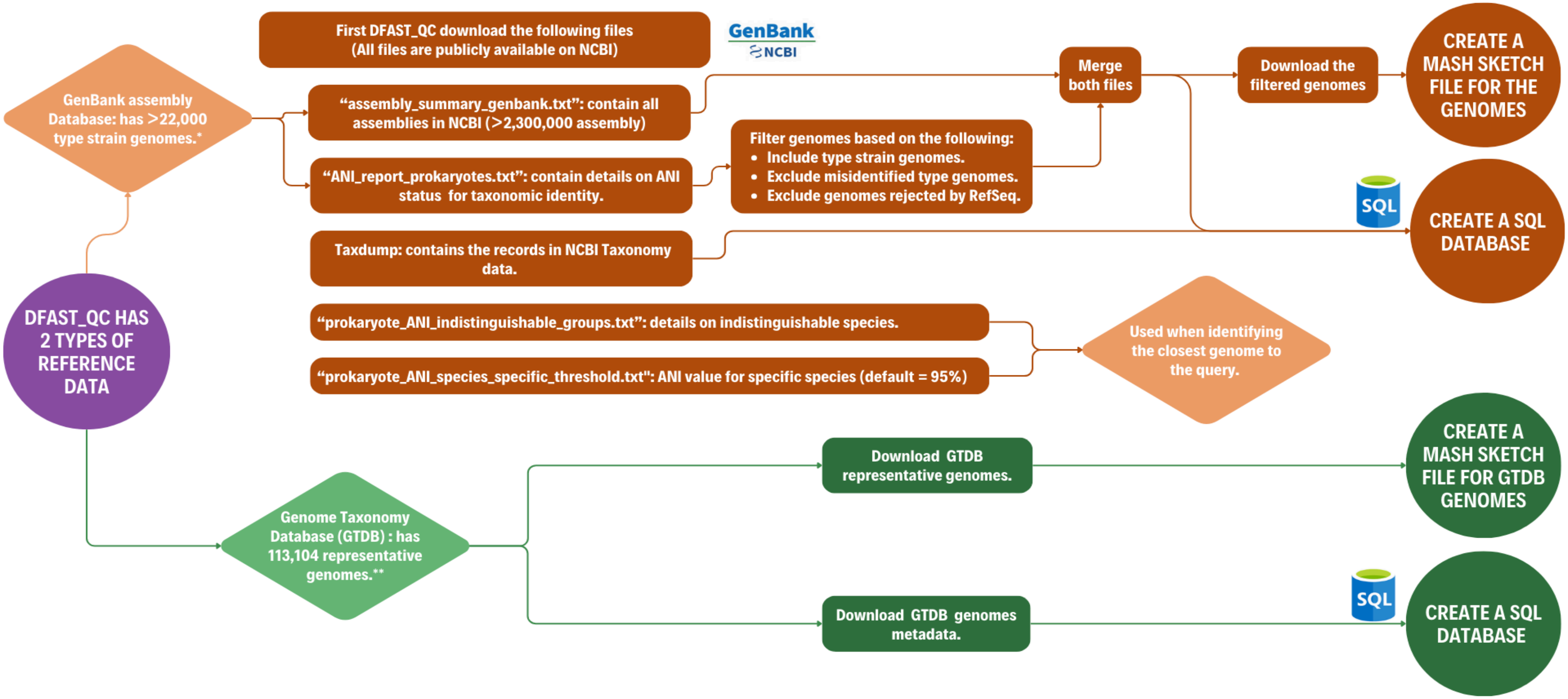

*As of June 26, 2024. **GTDB r220 release.
